# *An arch worth revisiting:* A study on the feline humeral supracondylar foramen and its evolutionary significance

**DOI:** 10.1101/2024.02.25.581957

**Authors:** Eimear Byrne, Robert D. Johnston, David Kilroy, Sourav Bhattacharjee

**Affiliations:** School of Veterinary Medicine, University College Dublin, Belfield, Dublin, Ireland; Trinity Centre for Biomedical Engineering, Trinity College Dublin, Dublin, Ireland; Department of Mechanical, Manufacturing & Biomedical Engineering, School of Engineering, Trinity College Dublin, Dublin, Ireland

**Author notes:** Corresponding author, T.

**Keywords:** supracondylar foramen, ligament of Struthers, median nerve, brachial artery, coracobrachialis longus, micro-computed tomography

## Abstract

The supracondylar foramen with a seemingly osseous peripheral arch noticed on the medio-distal feline humeri had remained disputed among anatomists. Some scholars have argued in favor of homology between this foramen and the supracondyloid foramen formed in the presence of the ligament of Struthers in humans. Other theories include its presence as a retinaculum holding the median nerve and brachial artery to their anatomical position in a flexed elbow. Unfortunately, these theories lack investigative rigor. The emergence of non-invasive imaging modalities, such as micro-computed tomography, has enabled researchers to inspect the internal anatomy of bones without dismantling. Thus, a micro-computed tomographic investigation was conducted on three feline humeri specimens while the internal anatomy of the supracondylar foramina was examined. Unlike the humerus, the thin peripheral arch of the feline supracondylar foramen failed to elicit any osseous trabeculae or foci of calcification. While adhering to the humeral periosteum at its origin, the non-osseous arch, typical of a muscular tendon or a ligament, inserts into a bony spur attached to the medial humeral epicondyle suggestive of a ligament or aponeurotic extension of a (vestigial) brachial muscle, with the coracobrachialis longus emerging to be the most likely candidate.

## 1. Introduction

The humeral supracondylar foramen (Figure 1), situated at the medio-distal aspect of the feline humeri—a few cm proximal to the medial epicondyle (Sánchez et al., 2013)—has intrigued anatomists for long. Although not ubiquitous, the presence of this foramen is not rare and is also noted in climbing mammals (e.g., lemurs) and marsupials (e.g., wombats and koalas). The structures that pass through this foramen are the brachial artery and median nerve.

**Figure 1.**
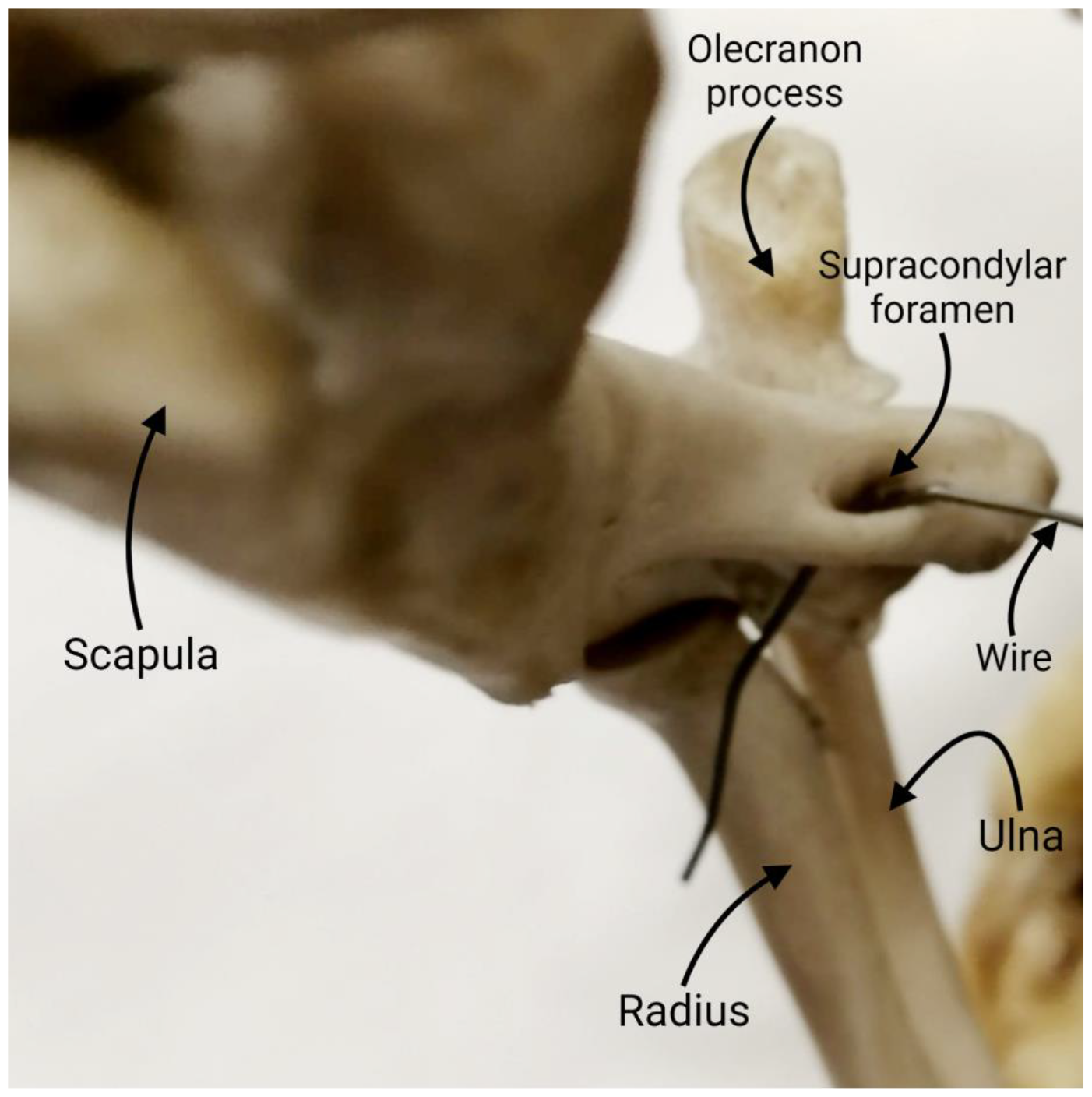
An oblique view of the elbow joint of a wallaby showing the articulating bones and the humeral supracondylar foramen (marked with a passage wire).

The function of this foramen remains obscure—and, unfortunately, fosters speculation. Some have argued that it plays a protective role for the neurovascular bundle of the forelimb during contraction of the overlying flexor muscles (Huntington, 1918), external blows, and fluctuating pressure felt, especially during abduction when the superficial structures become more exposed—thus are vulnerable to trauma (Tiedmann, 1822). On the contrary, some researchers have opined that the foramen or its apparently rigid peripheral arch signifies the attachment site for the pronator teres muscle with a shifted site of origin (Ruge, 1884), although this thesis was later contested (Stromer, 1902). Additionally, it serves as a brace or retinaculum (Figure 2) to prevent the median nerve from sagging forward into the angle of the elbow during flexion (Landry, 1958).

**Figure 2.**
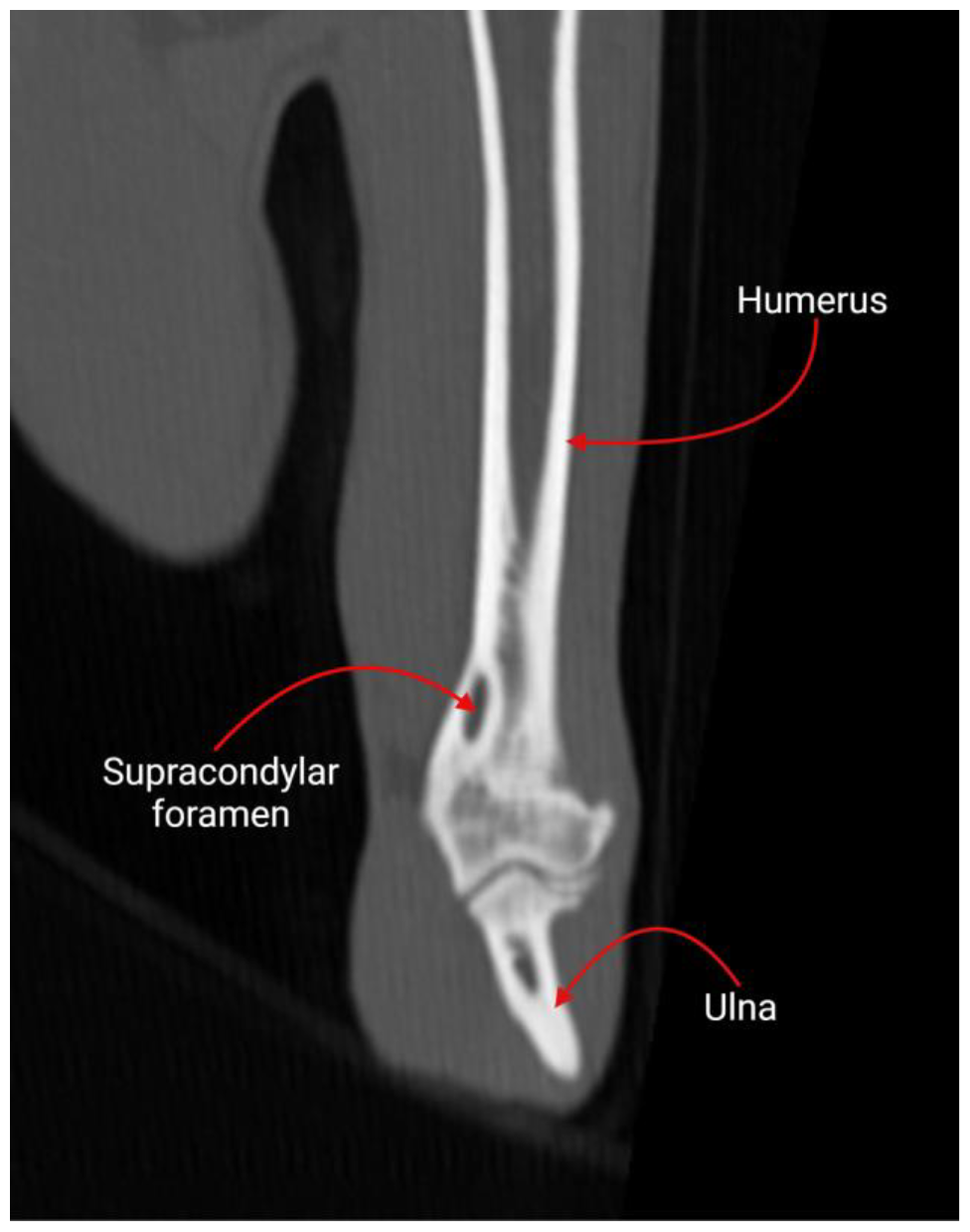
Computed tomographic image of a feline humerus showing the supracondylar foramen.

The filiform structures of the forelimb, including the vasculature, nerves, and muscle tendons, need to be held near the elbow joint during locomotion. The median nerve is especially vulnerable to such unwarranted mobility as it has no branches around the elbow to maintain its position. This exposes the neurovascular bundle to external trauma, given that it is only covered by overlying skin and a thin sheath of epitrochlearis muscle. Therefore, the theory of the supracondylar foramen acting as a retinaculum does bear merit. On the contrary, the brachial artery is relatively stable around the elbow joint due to its rich anastomoses and is not always encircled within the foramen, further strengthening the argument in favor of a protective role for the foramen.

The seemingly random occurrence of this foramen and its lack of consistency across species is also remarkable. It was suggested that the foramen is only present when the distal humeri are wide to allow a more direct route for the brachial artery and median nerve to access the antebrachium (Meckel, 1825). Conversely, a narrow humerus would obviate the need for such a foramen. However, this theory has lost traction as the foramen is obvious in feline species known to bear narrow distal humeri.

During an interspecies comparison, an interesting congruity was found between the feline species and a supposed homolog of the foramen in ∼2% of humans. Like the feline humeri, a bony spur, *viz*., supracondyloid process, is noted in the medio-distal aspect of human humeri a few cm proximal to the medial epicondyle (Natsis, 2008). This bony spur projects ∼2–20 mm from the medial humeral epicondyle and is connected to a thin ligamentous band of tissue, also known as the Struthers’ ligament—eponymized after the Scottish anatomist John Struthers (1823–1899)—that extends from the supracondyloid process to the medial epicondyle forming an arch over the median nerve and, occasionally, the brachial artery (Struthers, 1873). The arch is known to cause pain and paresthesia in the distal arm by causing compression to the median nerve (supracondylar process syndrome, Figure 3), often requiring surgical intervention (Opanova & Atkinson, 2014).

**Figure 3.**
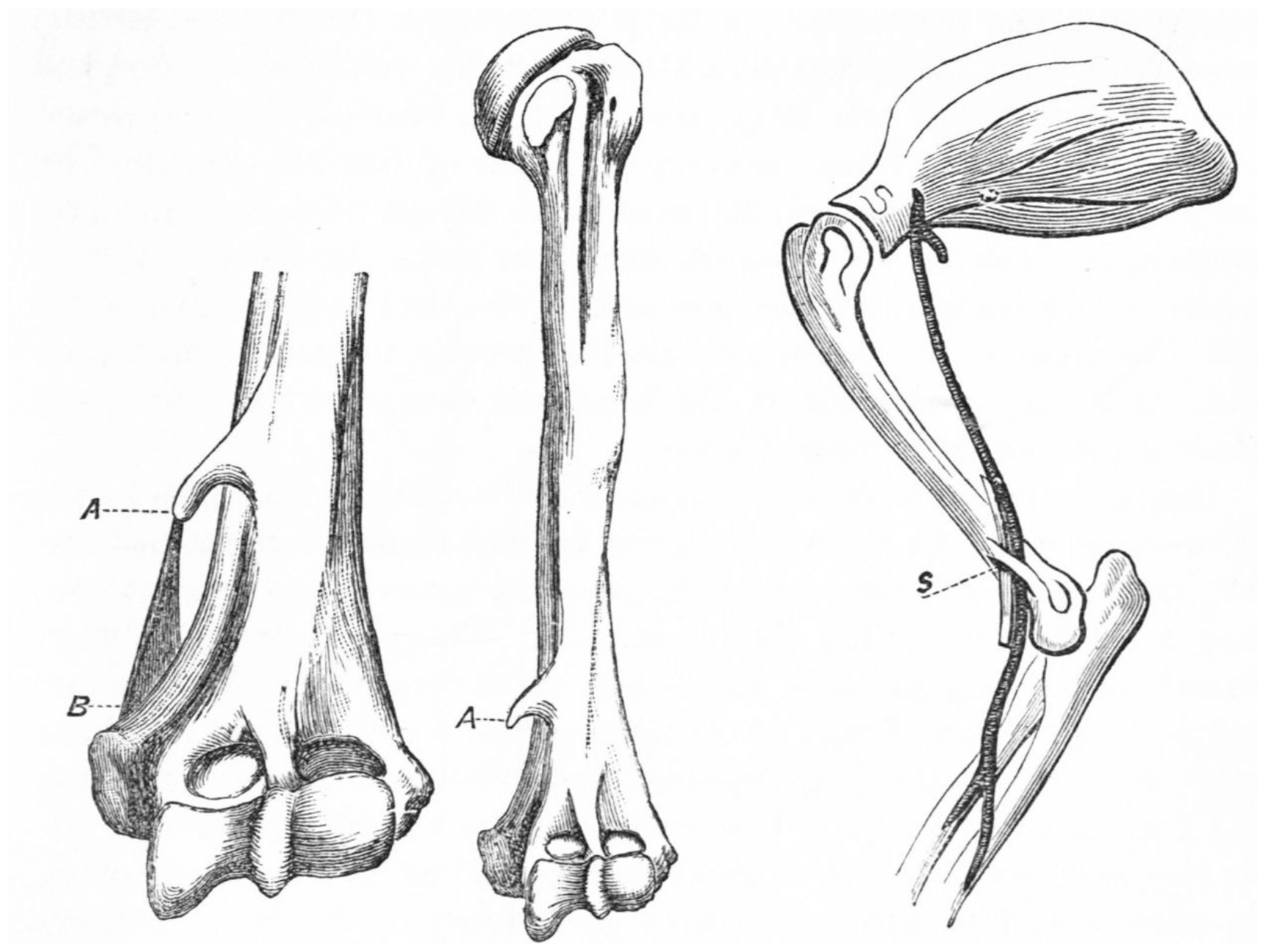
The analogy between the feline supracondylar ligament and the ligament of Struthers in humans. (A) The distally pointing supracondylar process proximal to the medial humeral epicondyle; (B) the ligament of Struthers as a fibrous band that, when present, forms the arch; (C) a medial view of the right feline humerus showing the supracondylar foramen with the median nerve and brachial artery passing through it. Modified and reproduced with permission from Struthers, 1873.

This study aimed to understand this peculiar supracondylar foramen, noticed in multiple species, including felines, from an evolutionary standpoint. Thus, an in-depth understanding of the internal fabric of the anatomical structures contributing to this foramen was deemed necessary. Our driving hypothesis is that this foramen, especially its peripheral edge, is not osseous—as it appears—but the remnant of a muscular tendon. Hence, non-invasive imaging tools, such as micro-computed tomography (micro-CT), were prioritized, given their ability to demonstrate granular details of anatomical structures without dismantling the specimens, such as bones (Hanson & Bagi, 2004).

Such emerging non-invasive tools provide imagery datasets that can further be analyzed with digital platforms, such as Fiji Is Just ImageJ (FIJI) software, an open-source image analysis platform that provides numerical readouts on their geometric, physical, and topographical attributes. The obtained data demonstrated the need to revisit such minute and peculiar anatomical structures, often considered serendipitous, that provide important insights into vertebrate evolution and, at times, warrant an upgrade—if not rectification—of our current understanding.

## 2. Materials and methods

### 2.1. Feline humeri

Three feline humeri specimens were collected from the anatomical specimen repository of the University College Dublin School of Veterinary Medicine and subjected to further micro-CT analyses. The study received an exemption from the Animal Research Ethics Committee of University College Dublin with an exemption code AREC-20-21-Kilroy.

### 2.2. Micro-CT data acquisition

The micro-CT on feline humeri was performed using a Nikon XT H 225 micro-CT scanner. The X-ray tube, equipped with the reflection 225 kV head, had the voltage set to 80 kV and a current of 140 μA, respectively. Furthermore, an aluminum filter of 0.5 mm thickness was used to filter the range of energies that interacted with the specimen. The stage position was then calibrated concerning the X-ray tube to ensure the specimen was within the field of view. In the acquisition stage, the field of view was set to capture the humeral supracondylar foramen with a voxel size of 20 μm. Ring correction and two averages were also performed to remove artifacts from the final images and ensure robust data collection. Scan time and reconstruction of the imaging data were approximately 12 h for the entire process. The acquired images were exported in JPEG format for further 3D surface projection in FIJI.

### 2.3. 3D surface projection in FIJI

FIJI is an open-source software freely available for download (https://fiji.sc/) and used for image computing, 3D rendition of (anatomical) structures, segmentation, and calculation of various 2D and 3D structural attributes for plotting and visualization. It is compatible with popular operating systems, including Windows, Linux, and macOS, and offers a user-friendly intuitive interface with a further opportunity to increase the depth of analyses by installing tailor-made plugins for precision.

For a 3D surface projection of the feline supracondylar foramen, the region of interest was cropped out of the micro-CT image followed by processing by widgets: FIJI → Analyze → 3D Surface Plot. The 3D surface plot was rendered in Thermal LUT (lookup table) under the following settings: Grid size: 256; Smoothing:12; Perspective: 0.2; Lighting: 0.69; z-scale: 0.22; Max: 55%; Min: 17%. The image scale was set using the following widgets: FIJI → Analyze → Set Scale.

## 3. Results

### 3.1. Internal features of the foramen

The micro-CT investigation acquired high-resolution scans through the feline humeri at different levels, and the *z*-stacks were then combined to develop a digital 3D rendition of the distal feline humerus in VG studio (https://www.volumegraphics.com/en/products/vgstudio.html) with a clear demonstration of its anatomical landmarks, including the trochlea and the supracondylar foramen (Figure 4A; Supplementary Video Files S1 and S2). The micro-CT data obtained from the other two feline humeri (Supplementary Material S3) corroborated with the representative specimen documented here. Internally, the bony trabeculae could be noticed while the humeral shaft was wrapped in a periosteum (Figure 4B). However, the typical osseous architecture was missing at the outer rim/arch of the foramen, and a shadow of less rigid, non-osseous tissue was detected (Figure 4B). This tissue, without traces of calcification, was attached to a well-defined bony spur attached to the medial condyle (Figure 4C).

**Figure 4.**
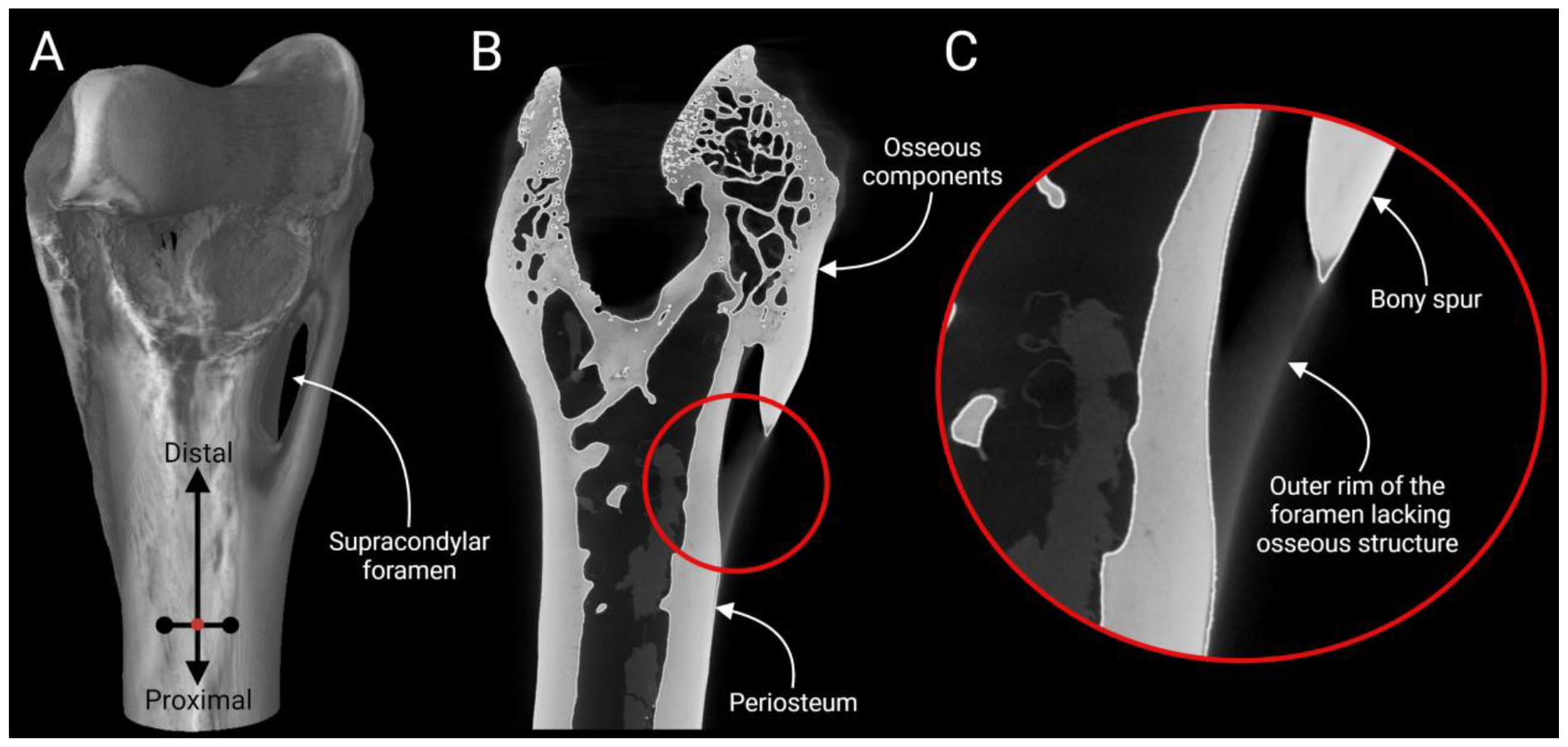
The distal feline humerus was viewed under micro-CT imaging. (A) Digital 3D reconstruction of the humerus; (B) the internal tissue fabric demonstrates a periosteum and trabeculae with a bony spur attached to a less rigid tissue; (C) a closer look into the bony spur with attached tissue band in a region of interest encircled within a red circle.

### 3.2. Osseous trabeculae

The micro-CT scan also revealed trails of extensive osseous trabeculae within the feline humerus, including its shaft, condyles, and articulating areas, whereas no such trabeculae could be identified in the unossified softer tissue mass connected to the medial condyle (Figure 5).

**Figure 5.**
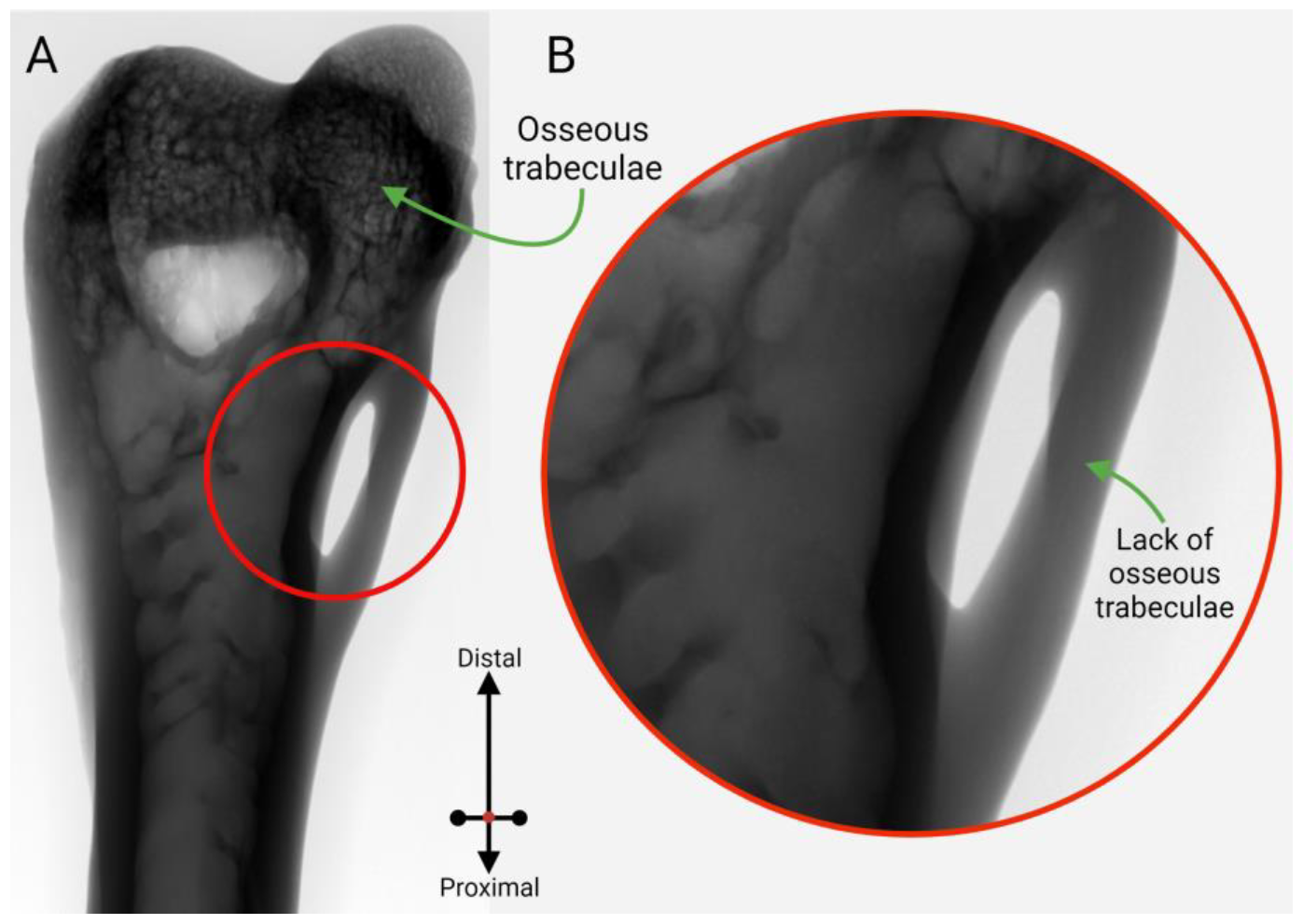
The scout view from a micro-CT scan demonstrated the osseous trabeculae trails inside a distal feline humerus. (A) Extensive trabeculae, characteristic of long bones, were noticed inside the humerus, including its shaft, condyles, and articulating surface, whereas no such traits were detected in the relatively soft unossified tissue forming the rigid outer arch of the foramen; (B) a closer look into the region of interest on the supracondylar foramen marked as a red circle.

### 3.3. 3D surface projection in FIJI

The 3D surface projection by FIJI categorically demonstrated the presence of a bony shaft attached to the edge of the humeral shaft. On the contrary, the tendinous remnant could be demarcated from the surrounding osseous structures, including the humeral shaft and the bony spur (Figure 6). The 3D rendition also demonstrated that the tendinous remnant was attached to the humeral shaft.

**Figure 6.**
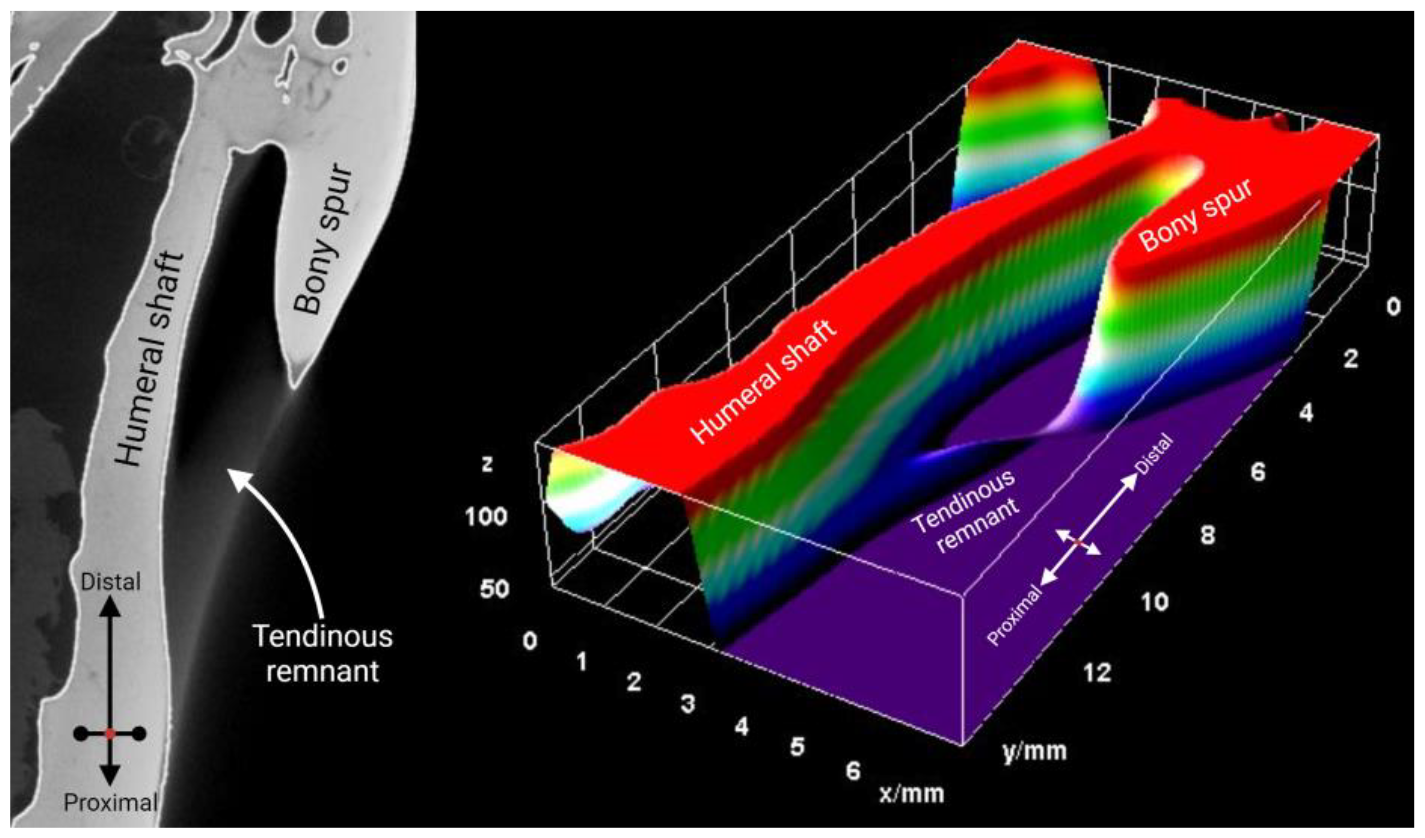
The 3D surface projection of a feline humeral supracondylar foramen rendered in FIJI software. The bony spur was attached to the humeral shaft, while the tendinous remnant, acting as the outer rim of the foramen, lacked an osseous fabric and was attached to the humeral shaft. The scale of the grids is included in the figure.

## 4. Discussion

The micro-CT investigation divulged granular details of the internal anatomy of the feline distal humeri. Moreover, with the help of *in silico* tools, such as FIJI, a realistic digital reconstruction of the feline humeri with crisp resolution could be achieved. The availability of such sophisticated imaging platforms in conjunction with advanced analyses harbors the potential of revolutionizing anatomy learning and research. It is a fair expectation that the utility landscape for such imaging modalities will keep expanding over time, and the research community will harness their benefits.

The micro-CT data have demonstrated the presence of a bony spur related to the medial humeral epicondyle and attached to an unossified band of tissue. It is worth noting here that there was speculation of it being a homolog of the supracondylar process attached to the ligament of Struthers (Kessel & Rang, 1966). Intriguingly, in his 1873 paper on the hereditary and humeral supracondyloid process in humans, John Struthers stated:

*“This process would, I believe, be found to occur as a variety also among some of the higher animals in which it does not exist normally, were they examined as often and as carefully as the human body is. I noticed it on the humerus of a fossil bear in Cuvier’s paleontological collection in the Jardin des Plantes, and I have the arm of a cat in which most of what should have been the bony archis represented by ligament. Someone will probably come upon the variety in one or other of the anthropoid apes as they increase in our collections*.*”*

Our data support this analogy, and an obvious similarity in external appearances further strengthens the thesis. It also demonstrates the power of modern imaging tools like micro-CT and data visualization in deciphering detailed anatomical structures. As confirmed by micro-CT, such a bony spur attached to a relatively softer tissue is characteristic of tendinous or ligamentous attachment at a muscular insertion, for example, at the insertion of the Achilles tendon on the calcaneal tuberosity (Pasetto et al., 2018). Under micro-CT, the softer tissues typically project hazy shadows compared to sclerotic osseous components due to a lack of calcification, and our data supported the notion that the supracondylar foramen’s arch is a remnant of a (vestigial) muscle.

This vestigial muscle may be a muscle of the arm (brachium). If true, it excludes a relation between the supracondylar and pronator teres, or in the case of cats, the epitrochlearis muscle, which are primarily muscles of the forearm(antebrachium). The coracobrachialis muscle is the only feasible brachial muscle contributing to this bony arch around the supracondylar foramen (Figure 7).

**Figure 7.**
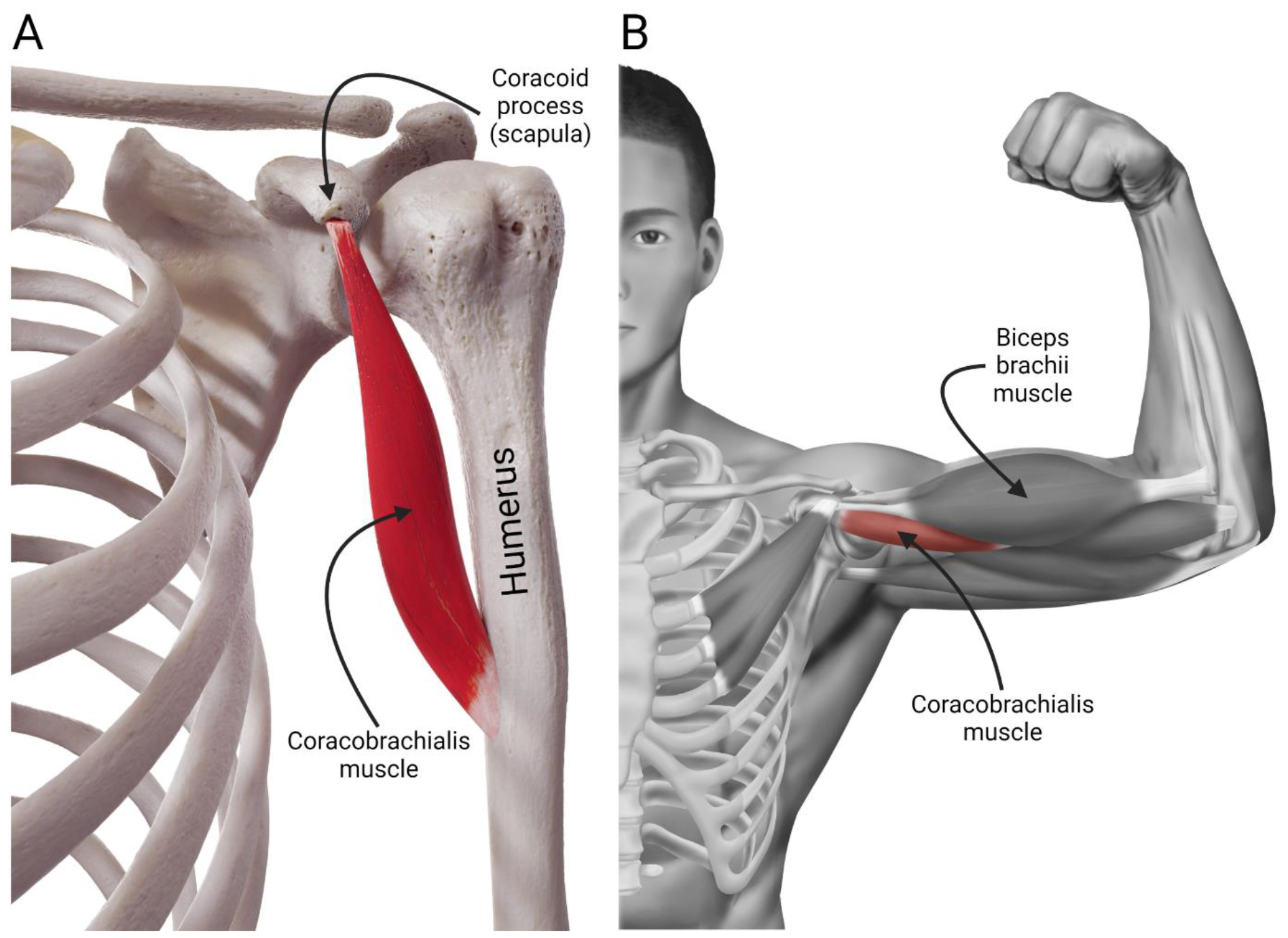
The anteroposterior views of the coracobrachialis muscle in humans, when the arm is in (A) anatomical and (B), abducted positions.

While taking its primary origin from the scapular coracoid process, the coracobrachialis muscle is inserted into the medial third of the humerus. It is the only surviving adductor muscle of the arm, and with hardly playing a functional role, it may be considered a vestigial muscle. In many species, including reptiles and amphibians, the muscle has three heads, *viz*., *coracobrachialis longus*—also known as Wood’s muscle—that is known to extend distally to the median nerve and brachial artery while staying related to the humeral shaft; *coracobrachialis medius* that is inserted into the humeral shaft distal to the insertion of latissimus dorsi muscle; and *coracobrachialis brevis* that inserts into medial humeral shaft proximal to the insertion of latissimus dorsi (Guha et al., 2010).

In humans, the coracobrachialis muscle has two heads surrounding the musculocutaneous nerve, and occasional nerve compression gives rise to clinical symptoms requiring surgical intervention. The coracobrachialis longus—although digressed almost entirely now—is known to give rise to a tendinous insertion on the humerus with an aponeurotic extension that reaches the distal end of the supracondylar ridge with ligamentous extension to the humeral inner condyle. Our data indicated that the unossified and softer tissue connected to the characteristic bony spur might have originated directly from the aponeurotic extension or its evolutionary remnant.

Perhaps the most important message distilled from our findings is the uncoupling of the supracondylar foramen from the ligament of Struthers. In the past, there had been plenty of confusion with the supracondylar foramen, ligament of Struthers, and the entepicondylar foramen—a foramen situated at the distal humerus of some amniotes—and noticed in fossils of archaic dinosaurs (Figueirido, 2020). Unfortunately, these terms have often been used injudiciously, and wild speculations made from purely external appearances further complicate comprehension.

This study has established that the supracondylar foramen, noticed in a feline skeleton, is a remnant from a vestigial arm muscle, with the coracobrachialis longus emerging as the most suitable candidate. However, an antebrachial muscle cannot be excluded completely. It is difficult to speculate which muscle it can be, although a proximally displaced origin of pronator teres emerges as a credible option, given that such a displacement might enjoy an evolutionary favor by providing improved grip to the forelimb due to better contraction. It is difficult to comment on the nature of entepicondylar foramen without micro-CT data, although, from external appearance, it seems more akin to the supracondylar foramen in cats.

As a historical note, even Charles Darwin was curious about such an opening near the medial humeral epicondyle and got in touch with John Struthers. In his book *The origin of man and selection in relation to sex* (1871), Darwin argued:

*“In some of the lower Quadrumana, in the Lemuridae and Carnivora, as well as in many marsupials, there is a passage near the lower end of the humerus, called the supra-condyloid foramen, through which the great nerve of the fore limb and often the great artery pass. Now, in the humerus of man, there is generally a trace of this passage, which is sometimes fairly well developed, being formed by a depending hook-like process of bone, completed by a band of ligament. Dr. Struthers, who has closely attended to the subject, has now shown that this peculiarity is sometimes inherited, as it has occurred in a father, and in no less than four out of his seven children. When present, the great nerve invariably passes through it; and this clearly indicates that it is the homologue and rudiment of the supra-condyloid foramen of the lower animals. Prof. Turner estimates, as he informs me, that it occurs in about one per cent of recent skeletons. But if the occasional development of this structure in man is, as seems probable, due to reversion, it is a return to a very ancient state of things, because in the higher Quadrumana it is absent*.*”*

The reported data here were able to discern between the feline supracondylar foramen and supracondylar process attached to the ligament of Struthers, with the possible muscle vestiges they may be assigned. The tetrapod evolution transitioned from water to land as their primary habitat during the Devonian period (419.2–358.9 Mya), and a fin-to-limb evolution of the forearm ensued. As a result, the distal end of the humerus has gone through considerable modulation. In sarcopterygian fishes, the median nerve and brachial artery run along a channel situated within a ventral crest on the humeral shaft before entering the forearm *via* a foramen, which can be taken as an archaic form of the entepicondylar foramen. However, with evolution, the course of the median nerve and brachial artery gradually changed, passing from the dorsal to the ventral side of the humerus (Landry, 1958).

This alteration in trajectories was perhaps accelerated by the emergence of *brachiation* that required restructuring of the brachial plexus, with a conversion of its directionality from an anteroposterior to a proximodistal fashion (Hirasawa & Kuratani, 2018) along with a drop and simultaneous torsion of the distal humerus resulting in depression. During the late Devonian and Carboniferous period (359.2–299 Mya), the entepicondylar foramen appeared to provide the median nerve and brachial artery a corridor to the forearm, ensuring a sustained—and perhaps an augmented—flow of blood while making concessions for a more extensive innervation plexus to cater to the need of increasing workload and complex biomechanics (Smithson & Clack, 2018).

Some have argued that the necessity of the entepicondylar foramen gradually diminished with the ungulates mostly moving their forelimbs in one plane without the need for humeral abduction. On the contrary, perhaps paradoxically, in humans, it has disappeared due to a much larger range of abduction (Landry, 1958). It is difficult to speculate on the evolutionary discourse of its appearance and subsequent disappearance—with occasional atavisms, such as the ligament of Struthers—in addition to its incredible variation noted even within the same species, family, or genus.

Weighing the data, it is prudent to accept that our understanding of the foramen near the medial humeral epicondyle needs a further boost. The complex genetic interactions and expressions should also be factored in. The data obtained in this study have elucidated that, at least in cats, the outer arch of the supracondylar foramen is a remnant from a vestigial muscle, which (possibly) is the coracobrachialis longus. However, the anatomy of the supracondylar process connected to the ligament of Struthers, or the entepicondylar foramen, remains far from settled.

An important lesson from this study is the relevance of advanced imaging tools in anatomical research, which perhaps remains the sole avenue available if the confusion, complexity, and the evolutionary trail between the feline supracondylar foramen, ligament of Struthers, and entepicondylar foramen are to be resolved. One of the reasons behind this confusion is that these three foramina are known for entrapping the same filiform structures of the forelimb: the median nerve and brachial artery. Moreover, their proximity to the medial epicondyle and an undeniable external similarity, especially in preserved specimens, leave ample room for dilemma and interpretation. That perhaps reiterates the primordial caution while dealing with anatomical specimens: *visual similarity does not guarantee equality*. The signatures of evolution, complex and minute, embedded in a tissue matrix, need revisitation through lenses that offer precision and resolution far ahead of what the human eyes can offer.

Given the extensive adaptation that the distal humerus underwent during its evolutionary journey, especially in its relations to the median nerve and brachial artery, any arch—ossified or not—near the medial humeral epicondyle will invariably engage these two structures, with full or partial entrapment while overarching.

Further follow-up investigations, perhaps facilitated with micro-CT, might reveal fascinating insights on entepicondylar foramen, indicating whether it is a mere homolog of these two structures mentioned earlier or stands alone as an independent anatomical landmark. Any such investigation on the entepicondylar foramen should also embrace old, fossilized specimens from primitive forms of vertebrates, dinosaurs included, as that may help raytracing its evolutionary trajectory and ambit. However, the success of micro-CT depends on the specimen material, and its quality, especially in fossilized specimens, unfortunately, will require plenty of optimization and data processing—to say the least—with a varying degree of reliability of the extracted output.

## 5. Conclusions

With the aid of micro-CT imaging, a study was conducted to investigate the internal tissue fabric of the feline humeral supracondylar foramen, which has remained an unsettled debate, especially from an evolutionary and morphological perspective. The obtained data demonstrated that, like ligaments or tendons, the outer arch of the foramen lacks calcification and bony trabeculae. Moreover, the relatively radiolucent arch attaches to a bony spur emerging out of the medial humeral epicondyle. The data supported the thesis of feline supracondylar foramen being a homolog or vestige to the arch formed by the presence of the ligament of Struthers anchored to a humeral supracondylar process. The obtained data indicated that the arch of this foramen is most likely a remnant of the tendon of the coracobrachialis longus muscle that, over time, has regressed and now remains a vestige. However, the exploits of antebrachial muscles like pronator teres cannot entirely be ruled out and need follow-up. The study showcased the power of advanced imaging tools like micro-CT in providing information on delicate structural details that, in alliance with emerging *in silico* data analysis and visualization suites, makes a strong case in favor of their augmented use in anatomical research and pedagogy.

## Acknowledgments

EB would like to thank the Anatomical Society for an Undergraduate Summer Vacation Research Scholarship (2020/21). EB, DK, and SB would like to thank Catherine McCarney from the UCD School of Veterinary Medicine for access to the feline humeri specimens and Michael Reilly from the Department of Mechanical, Manufacturing & Biomedical Engineering at Trinity College Dublin for arranging access to the micro-CT instrument.

The figures (except Figure 3) were created using Biorender.com.

## Author contributions

EB and RDJ conducted the research and data collection. DK and SB acquired funding, ethical exemption, and supervised the project. EB drafted the initial manuscript. All the authors have contributed to the final manuscript and approved the submitted version.

## Data availability statement

All the necessary data are reported within the manuscript. Further supportive data or clarifications are available from the corresponding author upon reasonable request.

## References

De Iuliis, G. & Pulerà, D. (2011) Chapter 7 - The cat. In: De Iuliis, G. & Pulerà, D. (eds.) The Dissection of Vertebrates (Second Edition). Boston: Academic Press, 147–252.

Figueirido, B. (2020) Vestiges of our first steps: An evolutionary view of the supracondylar syndrome. Mètode Science Studies Journal, 10, 213–220.

Guha, R., Satyanarayana, N., Reddy, C., Jayasri, N., Nitin, V., Praveen, G., Sunitha, P. & Datta, A. (2010) Variant insertion of coracobrachialis muscle-morphological significance, embryological basis and clinical importance. Journal of College of Medical Sciences-Nepal, (6)(2), 42–46.

Hanson, N. A. & Bagi, C. M. (2004) Alternative approach to assessment of bone quality using micro-computed tomography. Bone, 35, 326–333.

Hirasawa, T., & Kuratani, S. (2018) Evolution of the muscular system in tetrapod limbs. Zoological Letters, 4(1), 27.

Huntington, G. S. (1918). Modern problems of evolution, variation and inheritance in the anatomical part of the medical curriculum. The Anatomical Record, 14(6), 359–445.

Kessel, L. & Rang, M. (1966) Supracondylar spur of the humerus. The Journal of Bone and Joint Surgery. British volume, 48-B, 765–769.

Landry, S. O., Jr. (1958) The function of the entepicondylar foramen in mammals. The American Midland Naturalist, 60(1), 100–112.

Meckel, J. F. (1825) System der vergleichenden anatomie. Band 2, Abt. II. Leipzig: Halle.

Natsis, K. (2008) Supracondylar process of the humerus: Study on 375 Caucasian subjects in Cologne, Germany. Clinical Anatomy, 21, 138–141.

Opanova, M. I. & Atkinson, R. E. (2014) Supracondylar process syndrome: Case report and literature review. Journal of Hand Surgery, 39, 1130–1135.

Pasetto, L., Olivari, D., Nardo, G., et al. (2018) Micro-computed tomography for non-invasive evaluation of muscle atrophy in mouse models of disease. PLoS ONE, 13, e0198089.

Ruge, G. (1884) Beitrage zur Gefasslehre des Menschen. Morphologisches Jahrbuch, 9, 329–388.

Sánchez, H. L., Silva, L. B., Rafasquino, M. E., et al. (2013) Anatomical study of the forearm and hand nerves of the domestic cat (Felis catus), puma (Puma concolor) and jaguar (Panthera onca). Anatomia, Histologia, Embryologia, 42, 99–104.

Smithson, T. R., & Clack, J. A. (2018) A new tetrapod from Romer’s Gap reveals an early adaptation for walking. Earth and Environmental Science Transactions of the Royal Society of Edinburgh, 108(1), 89–97.

Stromer, E. (1902) Ueber die Bedeutung des Foramen entepicondyloideum und des Trochanter tertius der Saugethiere. Morphologisches Jahrbuch, 29, 553–562.

Struthers, J. (1873) On hereditary supra-condyloid process in man. The Lancet, 101, 231–232.

Tiedmann, F. (1822) Tabulae arterium corporis humani. Carlsruhae: Muller; 1822.

