## Supplementary Materials for "*An arch worth revisiting:* A study on the feline humeral supracondylar foramen and its evolutionary significance"

ORCiD

Eimear Byrne: https://orcid.org/0000-0002-9166-7841

Robert D. Johnston: https://orcid.org/0000-0002-2587-0606

David Kilroy: https://orcid.org/0000-0002-2692-8929

Sourav Bhattacharjee: https://orcid.org/0000-0002-6528-6877

*Corresponding author

T.: +353 1 716 6271

**Supplementary Material Figure S3.** The micro-CT data on the remaining two feline humeri specimens demonstrating the presence of bony spurs similar to the specimen reported in the main manuscript. A proximodistal arrow is embedded within the figure.

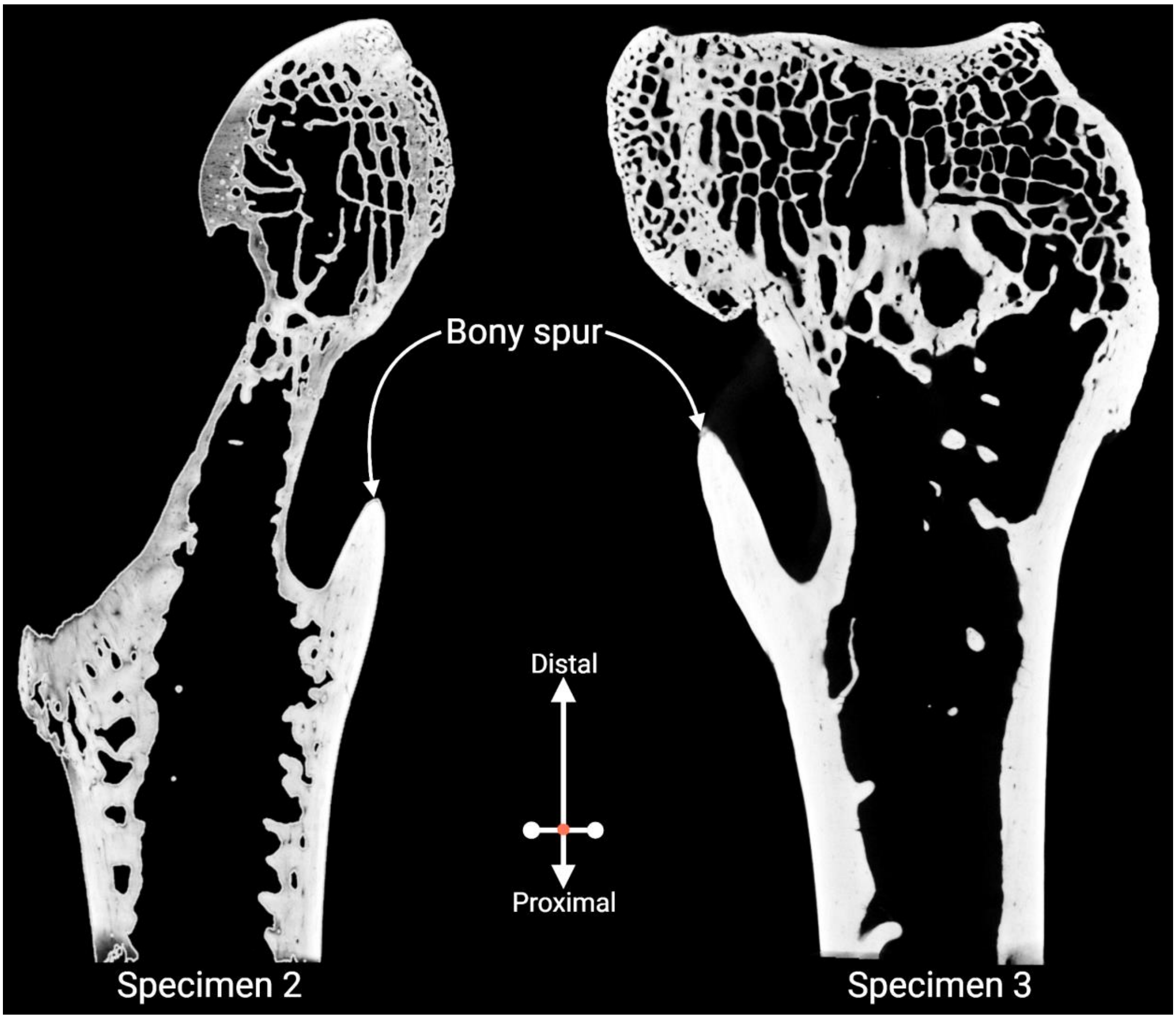
